# xml2jupyter: Mapping parameters between XML and Jupyter widgets

**DOI:** 10.1101/601211

**Authors:** Randy Heiland, Daniel Mishler, Tyler Zhang, Eric Bower, Paul Macklin

## Abstract

Jupyter Notebooks [4, 6] provide executable documents (in a variety of programming languages) that can be run in a web browser. When a notebook contains graphical widgets, it becomes an easy-to-use graphical user interface (GUI). Many scientific simulation packages use text-based configuration files to provide parameter values and run at the command line without a graphical interface. Manually editing these files to explore how different values affect a simulation can be burdensome for technical users, and impossible to use for those with other scientific backgrounds. xml2jupyter is a Python package that addresses these scientific bottlenecks. It provides a mapping between configuration files, formatted in the Extensible Markup Language (XML), and Jupyter widgets. Widgets are automatically generated from the XML file and these can, optionally, be incorporated into a larger GUI for a simulation package, and optionally hosted on cloud resources. Users modify parameter values via the widgets, and the values are written to the XML configuration file which is input to the simulation’s command-line interface. xml2jupyter has been tested using PhysiCell [1], an open source, agent-based simulator for biology, and it is being used by students for classroom and research projects. In addition, we use xml2jupyter to help create Jupyter GUIs for PhysiCell-related applications running on nanoHUB [5].

## Introduction

A PhysiCell configuration file defines model-specific *<*user_parameters*>* in XML. Each parameter element consists of its name with attributes, defining its data *type*, *units* (optional), *description* (optional), whether the widget should be *hidden* (optional), and the parameter’s default value. The attributes will determine the appearance and behavior of the Jupyter widget. For numeric widgets (the most common type for PhysiCell), xml2jupyter will calculate a delta step size as a function of the default value and this step size will be used by the widget’s graphical increment/decrement feature.

To illustrate, we show the following simple XML example, containing each of the four (currently) supported data types and the various attributes:

~~~
<PhysiCell_settings>
  <user_parameters>
    <radius type=“double” units=“micron”
      description=“initial tumor radius”>250.0
    </radius>
    <threads type=“int”>8</threads>
    <color type=“string” hidden=“true”>red</color>
    <fix_persistence type=“bool”>True</fix_persistence>
  </user_parameters>
</PhysiCell_settings>
~~~

When we map this into Jupyter widgets, we obtain the rendered results in Figure 1. Notice the color parameter is not displayed since we specified it should be hidden in the XML. The name of the other parameters, their values, and attributes, if present, are displayed in rows (as disabled Jupyter button widgets). Using alternating row colors (“zebra stripes”) helps visually match associated fields and avoid changing the wrong parameter value. For numeric widgets (type “int” or “double”), we compute a delta step value based on the magnitude (log) of the initial value. For example, the `radius` widget will have a step value of 10, whereas `threads` will have a step value of 1.

**Figure 1.**
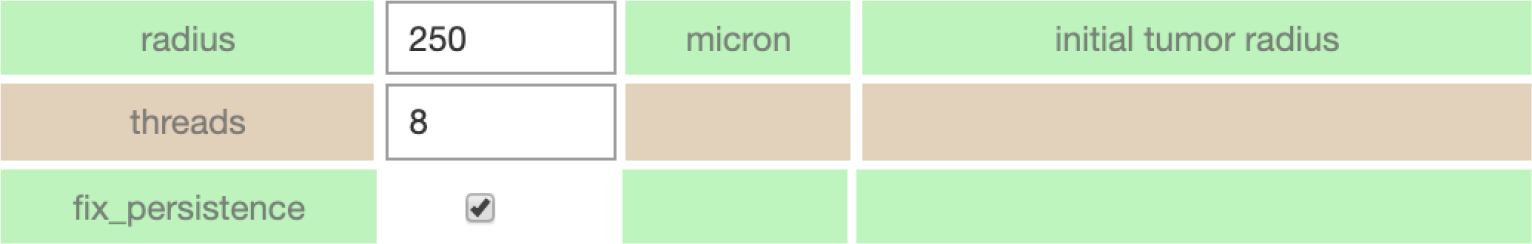
Simple example of XML parameters as Jupyter widgets.

For a more realistic example, consider the `config_biorobots.xml` configuration file (found in the `config_samples` directory). The XML elements in the *<*user_parameters*>* block include the (optional) *description* attribute which briefly describes the parameter and is displayed in another widget. To demonstrate xml2jupyter on this XML file, one would: 1) clone or download the repository, 2) copy the XML configuration file to the root directory, and 3) run the `xml2jupyter.py` script, providing the XML file as an argument.

~~~
$ cp config_samples/config_biorobots.xml.
$ python xml2jupyter.py config_biorobots.xml
~~~

The `xml2jupyter.py` script parses the XML and generates a Python module, `user_params.py`, containing the Jupyter widgets, together with methods to populate their values from the XML and write their values back to the XML. To “validate” the widgets were generated correctly, one could, minimally, open `user_params.py` in an editor and inspect it.

But to actually see the widgets rendered in a notebook, we provide a simple test:

~~~
$ python xml2jupyter.py config_biorobots.xml test_user_params.py
$ jupyter notebook test_gui.ipynb
~~~

This should display a minimal notebook in your browser and, after selecting `Run all` in the `Cell` menu, you should see the notebook shown in Figure 2.

**Figure 2.**
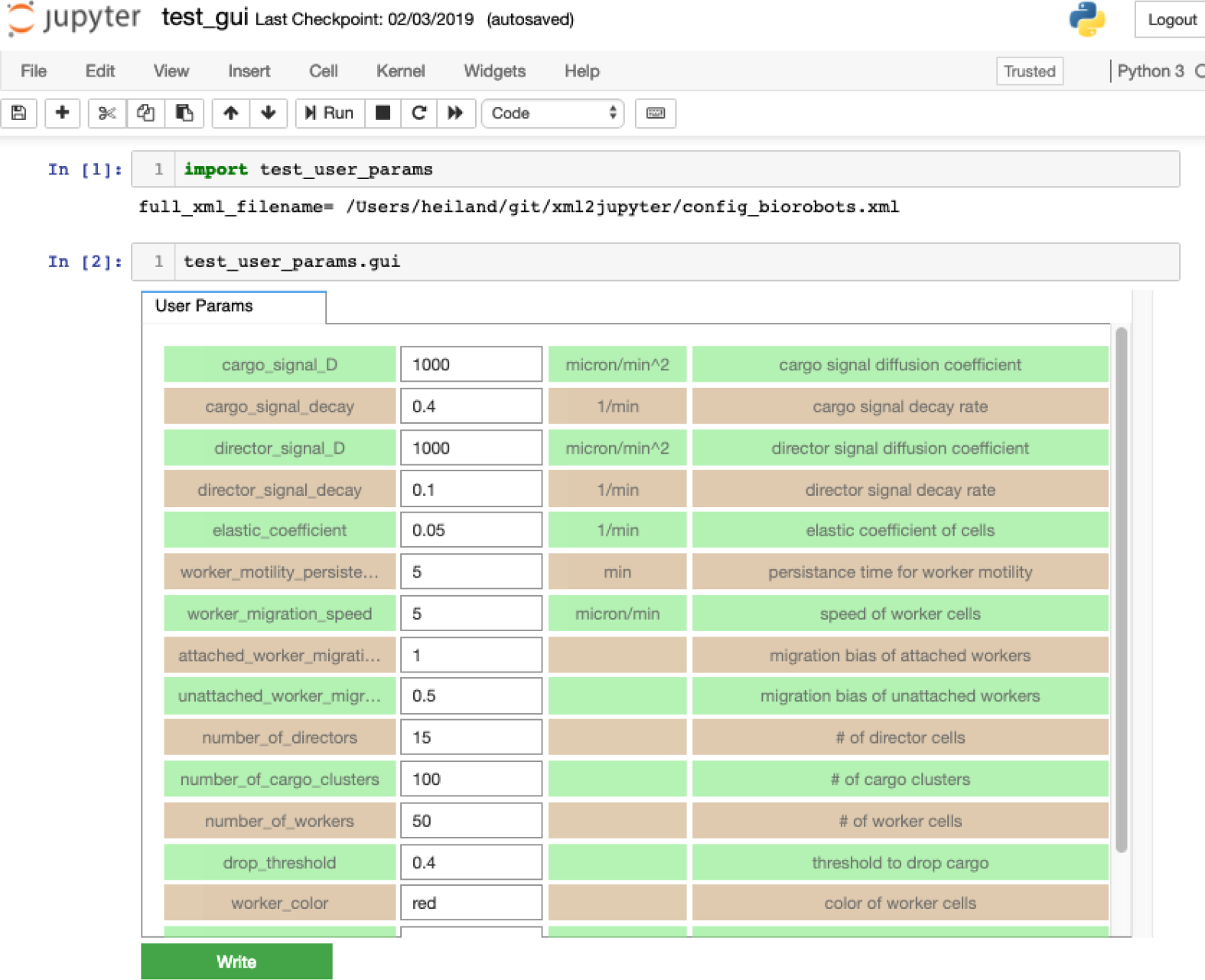
The biorobots parameters rendered as Jupyter widgets.

## PhysiCell Jupyter GUI

Our ultimate goal is to generate a fully functional GUI for PhysiCell users. xml2jupyter provides one important piece of this – dynamically generating widgets for custom user parameters for a model. By adding additional Python modules to provide additional components (tabs) of the GUI that are common to all PhysiCell models, a user can configure, run, and visualize output from a simulation. Two tabs that provide visualization of output files are shown below with results from the *biorobots* simulation. Note that some of the required modules are not available in the Python standard library, e.g., Matplotlib [2] and SciPy [3]. 2001). We provide instructions for installing these additional dependencies in the repository README.

## Extensions and Discussion

We hope others will be inspired to extend the core idea of this project to other text-based configuration files. XML is only one of several data-interchange formats, and while we created this tool for XML-based configurations based on needs to create GUIs for PhysiCell projects, the approach should be more broadly applicable to these other formats. And while the additional Python modules that provide visualization are also tailored to PhysiCell output, they can serve as templates for other scientific applications whose input and output file formats provide similar functionality.

xml2jupyter has helped us port PhysiCell-related Jupyter tools to nanoHUB, a scientific cloud for nanoscience education and research that includes running interactive simulations in a browser. For example, Figure 4 shows the xml2jupyter-generated *User Params* tab in our our `pc4cancerbots` tool running on nanoHUB. Figure 5 shows the cells (upper-left) and three different substrate plots for this same tool. This particular model and simulation is described in this video.

**Figure 3.**
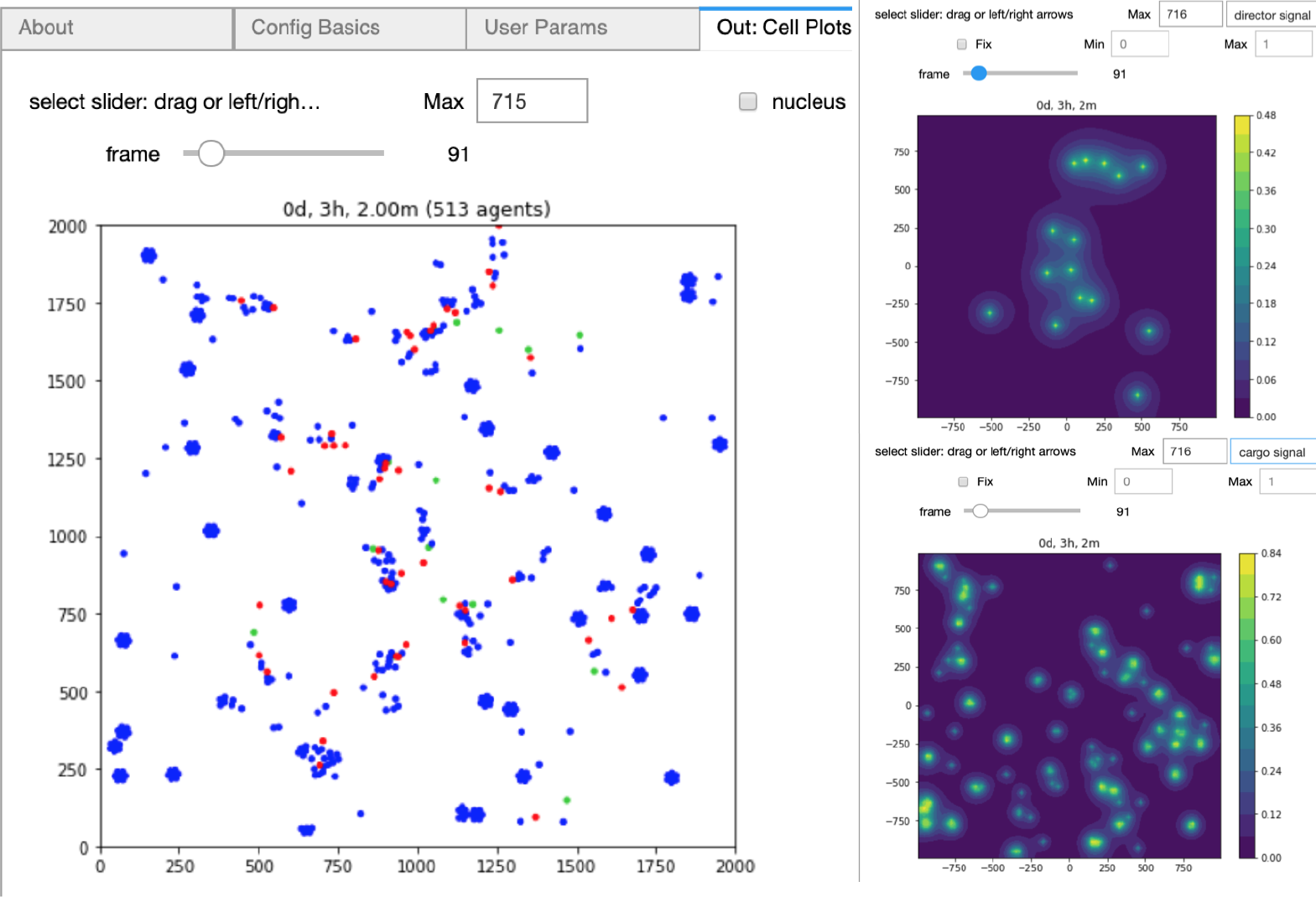
Plotting the biorobots (cells; left) and signals (substrates; right).

**Figure 4.**
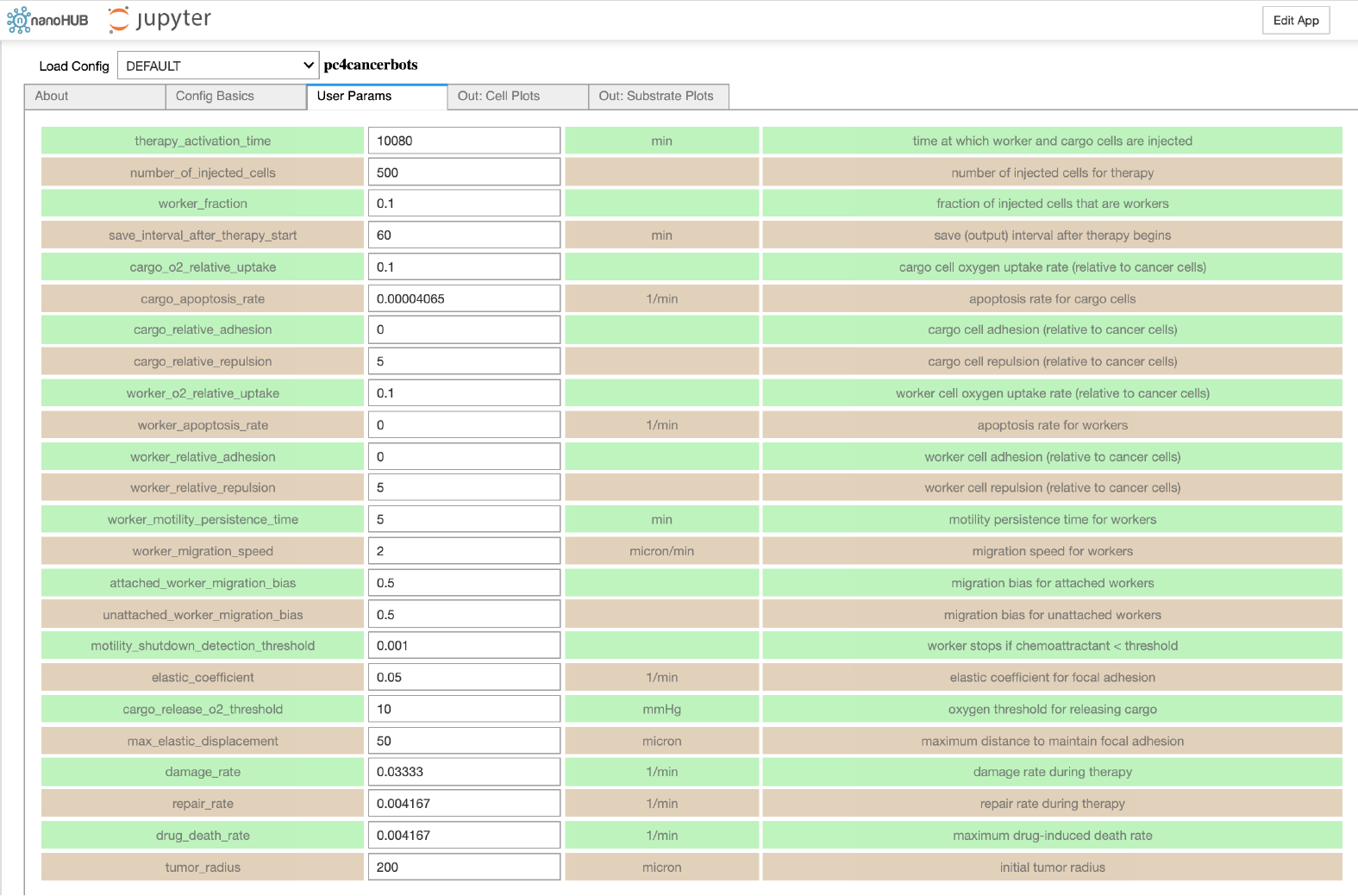
The cancer biorobots parameters in a nanoHUB Jupyter application.

**Figure 5.**
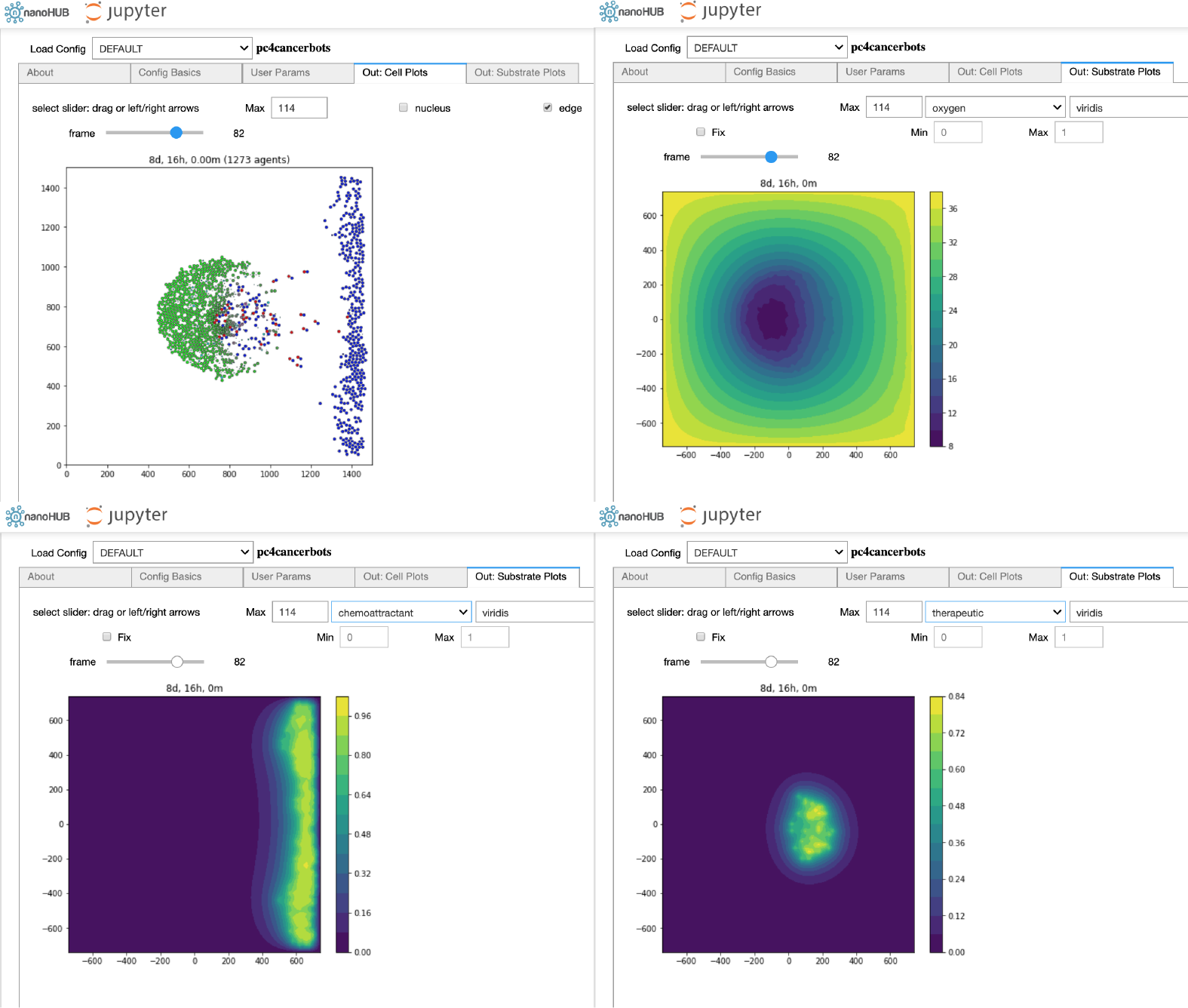
The cancer biorobots Jupyter notebook on nanoHUB.

Other PhysiCell-related nanoHUB tools that have been created using xml2jupyter include `pc4heterogen`, `pcISA`, and `pc4cancerimmune`. Readers can create a free account on nanoHUB and run these simulations for themselves. We encourage students to use xml2jupyter to create their own nanoHUB tools of PhysiCell models that 1) can be run and evaluated by the instructor, 2) can be shared with others, and 3) become part of a student’s living portfolio. (Another repository, https://github.com/rheiland/tool4nanobio, provides instructions and scripts to help generate a full GUI from an existing PhysiCell model.)

We welcome suggestions and contributions to xml2jupyter. For example, currently, we arrange the generated parameter widgets vertically, one row per parameter. This is an appropriate layout for an educational setting. But if a GUI will be used by researchers who are already familiar with the parameters, it may be preferable to generate a more compact layout of widgets, e.g., in a matrix with only the parameter names and values. Moreover, it may be useful to provide additional control over styling and placement by a separate style.xml or similar file, or by an external cascading style sheet. We will explore these options in the future, with the aim of separating as much GUI specification and styling from the original scientific application as possible. Such a decoupling would make it easier for scientific developers to continue refining their scientific codes without worrying about impact on the GUI, and without undue encumbrance by non-scientific annotations.

Also, we currently provide just 2-D visualizations of (spatial) data. In the near future, we will provide visualizations of 3-D models and welcome suggestions from the community.

## Acknowledgements

We thank the National Science Foundation (1720625) and the National Cancer Institute (U01-CA232137-01) for generous support. Undergraduate and graduate students in the Intelligent Systems Engineering deparment at Indiana University provided internal testing, and students and researchers within the NSF nanoMFG (1720701) group generously provided external testing. All of their feedback resulted in considerable improvements to this project. Finally, we thank our collaborators at Purdue University, especially Martin Hunt and Steve Clark, who provided technical support with nanoHUB and Jupyter.

